# *PIK3CA* Mutational Analysis in Formalin-Fixed, Paraffin-Embedded Archival Tissues of Urothelial Carcinoma of Urinary Bladder

**DOI:** 10.1101/020263

**Authors:** Alcides Chaux, Julie S. Cohen, Luciana Schultz, Roula Albadine, Sana Jadallah, Kathleen M. Murphy, Diana Taheri, Mark P. Schoenberg, George J. Netto

## Abstract

**Objective:** Urothelial carcinoma of the urinary bladder is the fourth most common cancer in males in the United States. In addition to mutations in *FGFR3*, *TP53*, *AKT1*, *TSC1*, and *PTEN* genes, mutations in *PIK3CA* have been also described in urothelial carcinomas, preferentially in low-grade tumors. Mutations in PIK3CA also has been shown to have implications for prognosis, surveillance and therapeutic response. Thus, determining the PIK3CA status in urothelial carcinomas could potentially improved the clinical management of patients with bladder cancer. Herein, we evaluated the presence of *PIK3CA* mutations in exons 1, 9, and 20 in 21 urothelial carcinomas of the urinary bladder.

**Methods:** Patients were treated by radical cystectomy without neoadjuvant chemotherapy. Representative tissue blocks (1 for each case) were selected. We used a pinpoint DNA extraction technique from formalin-fixed, paraffin-embedded and mutational analysis using the polymerase chain reaction (PCR) assay coupled with sequencing of targeted exons. Patients included 15 men and 6 women, with a median age of 68 years (range, 42 to 76 years), with 3 noninvasive and 18 invasive urothelial carcinomas. Noninvasive carcinomas included 1 case each of low-grade papillary urothelial carcinoma, high-grade papillary urothelial carcinoma, and urothelial carcinoma *in situ* (CIS). Invasive tumors included 3 pT1, 5 pT2, 6 pT3, and 4 pT4 urothelial carcinomas.

**Results:** We did not find mutations in the analyzed exons of the *PIK3CA* gene, in any of the 21 urothelial carcinomas. The preponderance of invasive high-grade and high-stage tumors could explain the absence of identifiable mutations in our cohort.

**Conclusions:** *PIK3CA* mutations as prognosticators of outcome or predictors of therapeutic response await further evaluation.

## 1 Introduction

Urothelial carcinoma of the urinary bladder is the fourth most common cancer in males in the United States, with an estimated 74,000 new cases and 16,000 deaths for 2015 [1]. The majority of the tumors are low grade, nonmuscle invasive tumors, associated with a good prognosis. However, approximately one-quarter of patients with bladder cancer are diagnosed with muscle-invasive tumors, with a significant risk of progression and a shortened survival [2]. It has been suggested that these two phenotypes of tumors progress through different pathways, which accounts for the differences in biological behavior. Low-grade, noninvasive tumors show high frequency in *FGFR3* mutation, whereas *TP53* mutations are associated with muscle-invasive tumors [3].

In addition to mutations in *AKT1*, *TSC1*, and *PTEN* [4, 5], mutations in *PIK3CA* have been also described in urothelial carcinomas, preferentially in low-grade tumors [6, 7, 8, 9, 10, 11, 12, 13, 14, 15]. Mutations in PIK3CA also has been shown to have implications for prognosis, surveillance and therapeutic response [9, 13, 16]. Thus, determining the PIK3CA status in urothelial carcinomas could potentially improved the clinical management of patients with bladder cancer. Herein, we evaluate the presence of *PIK3CA* mutations in exons 1, 9, and 20 in 21 patients with urothelial carcinomas of the urinary bladder. For this purpose, we use the polymerase chain reaction (PCR) coupled with sequencing of the targeted exons in formalin-fixed, paraffin-embedded tumor samples.

## 2 Material and Methods

The current study was approved by the Institutional Review Board at the Johns Hopkins School of Medicine (Baltimore, MD). The study has been performed in accordance with the ethical standards laid down in the 1964 Declaration of Helsinki.

### 2.1 Tissue Selection

Formalin-fixed, paraffin-embedded tissue samples of 21 patients with urothelial carcinoma of urinary bladder were selected from the pathology files of the Johns Hopkins Medical Institutions (Baltimore, MD). Patients were treated by radical cystectomy without neoadjuvant chemotherapy. Representative tissue blocks (1 for each case) were selected for microdissection and DNA extraction.

### 2.2 DNA Extraction from Formalin-Fixed, Paraffin-Embedded Tissue

Tumor areas were identified in routine sections stained with hematoxylin and eosin and 10 unstained sections (10 *μ*m thick) from each paraffin-embedded specimen were obtained. DNA isolation of the targeted tissue area on tissue sections was done using DNA Isolation System.

A drop of pinpoint solution (Pinpoint Slide DNA Isolation System; Zymo Research, Orange, CA) was applied to the mapped area of the tumor (approximately 5 × 5 mm^2^). Next, the targeted tumor tissue was microdissected with a scalpel and placed in a PCR tube. The excised tissues were digested in proteinase K buffer solution at 55 ° C for 8 hours, then at 97° C for 10 minutes.

### 2.3 *PIK3CA* Mutation Screening

PCR reactions were prepared with 1X PCR Buffer, 1.5 mM MgCl2, 500 *μ*M dNTPs (Applied Biosystems; Foster City, CA), 1.5 U AmpliTaq Gold (Applied Biosystems; Foster City, CA), and 500 nM each primer (Table 1) in a 50 *μ*l reaction. Six PCR reactions were performed to span the target regions: 2 covering exon 1; 1 covering exon 9; 3 covering exon 20 (Table 1).

**Table 1:**
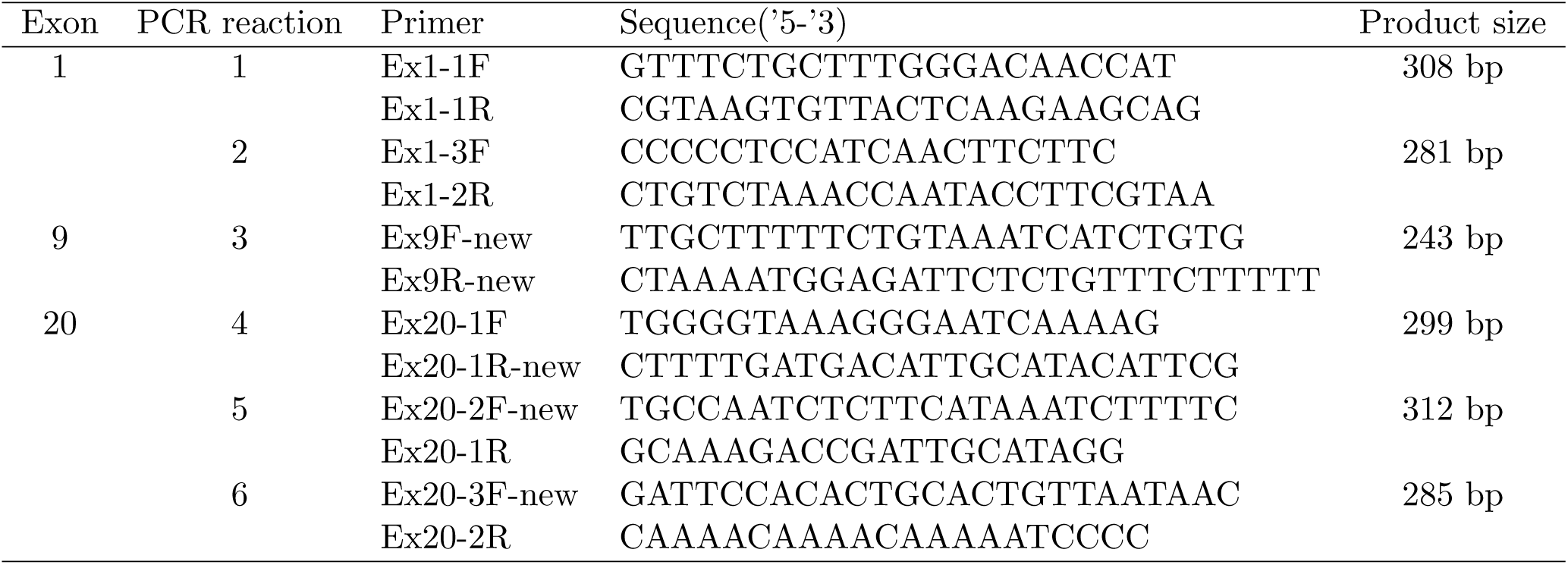
PCR Primers for PIK3CA Mutation Analysis

Reactions were heated to 95° C for 9 min followed by 35 cycles of 95° C for 30 sec, 53° C for 30 sec and 72° C for 1 min, followed by a final extension at 72o C for 7 min. Eight microliters of amplification product were separated by agarose (2%) gel electrophoresis to verify product. Amplification products were purified using QIAquick Spin Columns according to manufacturer’s protocol (Qiagen; Valencia, CA). Amplification products were cycle sequenced at an outside facility using Big Dye v3.1 reagents (Applied Biosystems; Foster City, CA).

Sequencing was performed in the forward and reverse directions using the PCR primers as sequencing primers (see Table 1). Sequencing products were purified using CleanSEQ Dye Terminator Removal reagents (Agencourt; Beverly, MA) and automated sequencing performed by capillary electrophoresis (CE) on an ABI3700 Avant genetic analyzer (Applied Biosystems; Foster City, CA). Sequence was analyzed with Sequencher 4.6 software (Gene Codes Corporation, Ann Arbor, MI). A cell line with *E545K* mutation was used as a positive control.

## 3 Results

### 3.1 Clinicopathologic Data

The group of patients was composed of 15 men and 6 women, with a median age of 68 years (range 42–76 years). Cases included 3 noninvasive and 18 invasive urothelial carcinomas. Noninvasive carcinomas corresponded to 1 case each of low-grade papillary urothelial carcinoma, high-grade papillary urothelial carcinoma, and “flat” *in situ* urothelial carcinoma. Invasive tumors included 3 pT1, 5 pT2, 6 pT3, and pT4 urothelial carcinomas.

### 3.2 *PIK3CA* Mutation Analysis of exons 1, 9, and 20

In 2 cases (1 pT2 and 1 pT4), PCR reactions were not informative for any of the analyzed exons. In 3 additional cases (1 pT1, 1 pT3, and 1 pT4), the PCR reaction was not informative for exon 9. In all the remaining cases and exons, no evidence of *PIK3CA* mutations was found.

## 4 Discussion

Herein, we evaluated the presence of *PIK3CA* mutations in urothelial carcinoma of the bladder using formalin-fixed, paraffin-embedded archival tissues. Entire exons 1, 9, and 20 were covered using the PCR assay coupled with sequencing of the targeted exons. No *PIK3CA* mutations were identified in any of the 21 patients under examination. Our results differ from previous studies, in which *PIK3CA* mutations were identified in 13% to 27% of bladder tumors [6, 7, 11, 12, 8]. Since most of our cases were highgrade urothelial carcinomas, this discrepancy might be due to the association between *PIK3CA* mutation and lower tumor grade, as suggested by previous studies [6, 7, 8] and discussed below.

*PIK3CA* mutations have been previously characterized in bladder cancer. Ló pez-Knowles *et al* sought for mutations in exons 9 and 20 of *PIK3CA* using paraffin-embedded tissue from 87 patients with bladder cancer [6]. Eleven (13%) tumors harbored *PIK3CA* mutations, and the prevalence was significantly higher in low-grade tumors. In the same study, authors reported *PIK3CA* mutations in 26% of 43 patients with papillary urothelial neoplasm of low malignant potential (PUNLMP). Similar results were found by Platt *et al* who identified *PIK3CA* mutations in 27% of 92 tissues samples [7]. They also observed a significant relationship with tumor grade but not with tumor stage.

Kompier *et al* evaluated 257 patients with bladder tumor and found *PIK3CA* mutations in 24% of the tissue samples [8]. Although the prevalence of *PIK3CA* mutations was higher in low grade tumors, there was no statistical significance regarding tumor grade. In another recent study, Sjödahl *et al* investigated the role of several genes in urothelial carcinomas, including *PIK3CA* and PIK3R1 [10]. *PIK3CA* was mutated in 37 of 218 patients (17%), and was associated with low grade tumors. In contrast to previous studies, higher proportion of tumors harboring *PIK3CA* mutation was seen in pTa tumors compared to pT1 tumors. However, the difference did not hold between pT1 and muscle-invasive (*>*pT1) tumors. Our results confirm the feasibility of *PIK3CA* mutational analysis in formalin-fixed, paraffin-embedded tissue samples and the rarity of *PIK3CA* mutations in high-grade and muscle-invasive urothelial carcinomas.

The usefulness of *PIK3CA* mutations as prognostic factors has been also explored. Lindgren *et al* classified urothelial carcinomas in 2 groups defined by gene expression, and found that *PIK3CA* mutations are significantly more frequent in the subtype of tumor that is related to a better prognosis [9]. Dueñas *et al* found that the presence of *PIK3CA* mutations is significantly associated with reduced recurrence in patients with non-muscle invasive bladder cancer [13]. Kim *et al* found that *PIK3CA* mutations are associated with improved recurrence-free survival and improved cancer-specific survival in patients with high-grade urothelial carcinoma of urinary bladder treated by radical cystectomy [16]. In spite of these aforementioned studies, Kompier *et al* found that *PIK3CA* mutations are not independent predictors of tumor recurrence, tumor progression, or disease specific survival [8]. Nevertheless, in the study by Kompier *et al*, patients with recurrence showed a 100% concordance in the type of *PIK3CA* mutation between tumor samples. This finding suggests a potential utility for *PIK3CA* in the screening and follow-up of patients with bladder tumors harboring such mutation. Another direction for future studies could be the putative impact of *PIK3CA* mutations in the therapeutic response of patients with superficially-invasive urothelial carcinomas treated locally.

In summary, we analyzed 21 formalin-fixed, paraffin-embedded samples from patients with urothelial carcinomas of the urinary bladder, seeking to identify *PIK3CA* mutations. Exons 1, 9, and 20 were fully covered using a pinpoint DNA extraction technique and PCR assay. We did not find any mutations in the *PIK3CA* gene. Most of our cases were high-grade and/or high-stage tumors, which could explain the absence of identifiable mutations. The role of *PIK3CA* mutations as prognosticators of outcome or predictors of therapeutic response awaits further evaluation.

## 5 Financial Disclosure

This study was supported by The Johns Hopkins Medicine - Patana Fund for Research, PO1# CA077664 NCI/NIH Grant, David H. Koch Prostate Cancer Fund, and the Flight Attendant Medical Research Institute (FAMRI) Clinical Innovator Award. Dr. Alcides Chaux was partially supported by an award granted by the National Council of Science and Technology (CONACYT) dependent of the Presidency of the Republic of Paraguay, as an Active Researcher of Level 2 of the National Incentive Program for Researchers (PRONII).

